# Buying time: detecting Vocs in SARS-CoV-2 via co-evolutionary signals

**DOI:** 10.1101/2022.07.21.500897

**Authors:** Christopher Barrett, Andrei C. Bura, Qijun He, Fenix W. Huang, Thomas J. X. Li, Christian M. Reidys

**Affiliations:** Biocomplexity Institute and Initiative, University of Virginia, Charlottesville, VA, USA; Department of Computer Science, University of Virginia, Charlottesville, VA, USA; Department of Mathematics, University of Virginia, Charlottesville, VA, USA

**Keywords:** motif-complex, co-evolution, SARS-CoV-2, VOC alert

## Abstract

We present a novel framework facilitating the rapid detection of variants of interest (VOI) and concern (VOC) in a viral multiple sequence alignment (MSA). The framework is purely based on the genomic sequence data, without requiring prior established biological analysis. The framework’s building blocks are sets of co-evolving sites (motifs), identified via co-evolutionary signals within the MSA. Motifs form a weighted simplicial complex, whose vertices are sites that satisfy a certain nucleotide diversity. Higher dimensional simplices are constructed using distances quantifying the co-evolutionary coupling of pairs and in the context of our method maximal motifs manifest as clusters. The framework triggers an alert via a cluster with a significant fraction of newly emerging polymorphic sites. We apply our method to SARS-CoV-2, analyzing all alerts issued from November 2020 through August 2021 with weekly resolution for England, USA, India and South America. Within a week at most a handful of alerts, each of which involving on the order of 10 sites are triggered. Cross referencing alerts with a posteriori knowledge of VOI/VOC-designations and lineages, motif-induced alerts detect VOIs/VOCs rapidly, typically weeks earlier than current methods. We show how motifs provide insight into the organization of the characteristic mutations of a VOI/VOC, organizing them as co-evolving blocks. Finally we study the dependency of the motif reconstruction on metric and clustering method and provide the receiver operating characteristic (ROC) of our alert criterion.

## Introduction

Current genomic sequencing efforts facilitate virological epidemiological surveillance close to real time. The challenge is to efficiently identify variants within these viral sequences that pose further threats. We present here a bottom up framework facilitating the rapid detection of variants of interest (VOI) and of concern (VOC), given a times series of multiple sequence alignments (MSA) consisting of viral genomes. The key idea is to identify maximal sets of sites exhibiting co-evolutionary signal within the MSA instead of considering the emergence of particular mutations. These signals naturally induce a complex of motifs formed by sets of co-evolving sites. The sites are selected on the basis of exhibiting sufficient mutational activity and satisfy a certain diversity criterion. Higher dimensional simplices are constructed using distances capturing the co-evolutionary coupling of pairs.

Via this method we develop and analyze an alert protocol: an alert is triggered by a cluster exhibiting a significant fraction of newly emerging sites. Then our alerts are put to the test, analyzing retrospectively SARS-CoV-2 sequence data collected from November 2020 through August 2021. That is we issue alerts based on “historic” data, not “real-time” alerts. These alerts are issued with no *a priori* assumptions, except of the Wuhan reference sequence upon which the MSAs are built. MSAs are constructed on a weekly basis for England, USA, India and South America (SA). Thereafter we relate our alerts to established VOIs and VOCs, i.e. employing the *a posteriori* knowledge of VOI/VOC-designations and lineages in order to evaluate the accuracy of the issued alerts. We remark that an alert does not provide any biological semantics, its primary purpose is to enable a fast biological analysis on a handful of critical sites.

### Background

Genomic surveillance plays an instrumental role in combating rapidly mutating RNA viruses [14]. In particular, it is becoming a vital necessity in the effective mitigation and containment of the COVID-19 pandemic [41, 8]. While mRNA vaccine development and distribution was successful in the US, recently VOCs, in particular the Delta-variant [32], give rise to questions regarding the efficacy of current vaccines and timely vaccine development necessitates the rapid recognition of critical adaptations within SARS-CoV-2. Genomic surveillance leverages applications of next-generation sequencing and phylogenetic methods to detect variants that are phenotypically or antigenetically different, facilitating early anticipation and effective mitigation of potential viral outbreaks.

One of the central tasks in genomic surveillance is to identify emerging variants that are more virulent, or more resistant to available vaccines. The designation of SARS-CoV-2 variants of concern/interest exemplifies such an identification process [26, 48]. Currently, the designation of such a VOC is based on phylogenetic methods and involves four steps: lineage assignment, mutation extraction, biological analysis, and declaration. First, a large phylogenetic tree is constructed from publicly available SARS-CoV-2 genomes, and its sub-trees are examined and cross-referenced against epidemiological information to designate new lineages [3, 40]. Secondly, a collection of mutations, frequently observed in a lineage is extracted and defined to be characteristic. The biological impact of this collection of mutations is then analyzed in wet-lab/in silico experiments. Thirdly, in the wake of identified biological features, such as an increase in transmissibility or severity, the lineage/variant is declared a VOC.

Population-based approaches were developed to complement phylogenetic-based methods with the goal of rapidly identifying and monitoring critical mutations on the SARS-CoV-2 genome. Frequency analysis is widely used to monitor variant circulation [27, 35]. The increasing prevalence of a mutation might indicate the emergence of a new variant. Entropy measurements, derived from nucleotide frequency, highlight nucleotide positions with high variation and facilitate the compact representation of SARS-CoV-2 variants [12].

Mutations on viral genomes do not always appear independently. For example, the D614G (A23404G) mutation on the SARS-CoV-2 genome is almost always accompanied by three other mutations: C241T, C3037T and C14408T [37]: these four positions exhibit a co-evolutionary pattern. In fact, positions in a molecule that share a common constraint do not evolve independently, and therefore leave a signature in patterns of homologous sequences [11, 38]. Extracting such co-evolution signals from a sequence alignment leads to a deeper understanding of the impact of mutations and can facilitate the early detection of emerging variants. Present correlation analysis techniques are not amenable to co-evolutionary analysis. For instance, the Pearson correlation coefficient [36] requires the computation of the average value of a random variable. While the nucleotide type at a fixed position in an alignment can be regarded as a random variable, averaging techniques are not straightforwardly applicable. While Spearman’s correlation coefficient [34] and the Kendall Tau rank coefficient [25] work for rank correlation analysis, there exists no canonical ranking for the different types of nucleotides. Co-evolution detection strategies currently are based on observing the frequency of nucleotide combinations in two distinguished positions [31, 47]. It is a challenge to dynamically keep track of all such pairwise frequencies on sizable data sets. Thus, a novel measurement for the degree of co-evolution is of relevance.

In protein folding MSA are employed to identify related positions [46] via mutual information. While there are contributions on the level of networks: [1] studies mutual information networks of enzymatic families in protein structures to unveil functional features, the work is focused on how to account for the effect of phylogeny on this identification [6]. To this end, two modifications to mutual information are introduced: row-column weighting [15] and average product correction [10]. Mutual information has also been used to detect coevolution signals in alignments of RNA sequences [17]. In [16, 49] statistical methods differentiate correlation patterns induced by functional constraints from those induced by shared ancestry. These methods are concerned with pairwise relations since their objective is to determine RNA secondary structure.

Thus, the idea of considering pairwise relations between columns within an MSA has been successfully used in protein and RNA folding more than a decade ago. The motif complex represents an extension of this, encapsulating *k*-ary relations within the MSA, that cannot be reduced to pairwise relations. When applying our method to SARS-CoV-2, however, we consider only pairwise relations. This allows us to use mutual information and standard clustering algorithms, where clusters approximate maximal motifs. The motif complex suggests extending the notions of mutual information and distance beyond two random variables and points, respectively.

### Motifs and alerts

The framework developed here represents a bottom-up approach requiring no *a priori* knowledge of phylogeny, lineages or any type of biological impact analysis. Its output consists of a collection of a small number of distinguished clusters composed by critical, tightly co-evolving positions on the SARS-CoV-2 genome. The particular nucleotide identity at these positions, as relevant as they are for subsequent analysis, only plays a subordinate role. The notion of a reference sequence also plays a substantially different role: it is exclusively employed for the generation of the multiple sequence alignment from which the aforementioned notion of position/site (i.e. column in the alignment matrix) originates. A group of mutations co-evolving via a similar pattern exhibits footprints of evolutionary selection pressure. The fitness induced by a group of co-evolving mutations can be more significant than their total fitness when they occur independently, as hidden links might exist between the sites in question. Therefore, a group of sites having sufficient nucleotide diversity that are clustered by means of co-evolution measures can represent a signal of selective advantages.

We consider here clusters that carry a significant portion of newly active sites. These sites represent a sufficiently large additive fitness component of the underlying cluster and are potentially indicative of the emergence of a functional block, namely a keystone mutation event. The alert picks up the induced differential in the evolutionary dynamics, very much in the spirit of a derivative. Specifically, alerts (Section *The motif complex of SARS-CoV-2*) are closely related to the derivate of the logarithm of the size of the cluster inducing the alert. We are in effect constructing a guidance system, alerting not only to the sites where biological analysis should be performed, but also quantifying at which rate they co-evolve. This provides crucial information for biologists, since identifying co-evolution relations provides clues about underlying biological mechanisms.

GISAID-sequence data are rich enough to allow for a weekly time resolution and within this timeframe towards a handful of alerts, each involving on the order of 10 sites are triggered. Relating alerts with the *a posteriori* knowledge of VOI/VOC-designations and lineages, motif-induced alerts detect VOIs/VOCs rapidly, typically weeks earlier than current methods. We show how motifs provide insight into the organization of the characteristic mutations of a VOI/VOC, organizing them as co-evolving blocks. Finally we study the dependency of the motif reconstruction on metric and clustering method and provide the receiver operating characteristic (ROC) of the alert criterion.

## The motif complex

In this section we specify a mathematical framework that allows us to express co-evolutionary signals in MSA. In a natural way these signals give rise to a weighted, simplicial complex, upon which our notion of alert derives.

Let *A* = (*a*_*p,q*_) denote a multiple sequence alignment (MSA) composed of *m* sequences of length *n*. Here *a*_*p,q*_ *∈* 𝔸 = {*e, A, T, C, G*} represents the nucleotide at the *q*th position of the *p*th sequence and *e* denotes a gap.

We consider a *k*-tuple of *A*-columns *i*_1_, *i*_2_, …, *i*_*k*_ and query the existence of a *k*-ary relation between the nucleotides present at *i*_1_, *i*_2_, …, *i*_*k*_, respectively. We stipulate that such a relation is the result of selective pressure exerted on a collective of sites forming a block having implicit connection in the viral genome. Such a site-dependency can manifest via a variety of mutational constellations.

For each collection of sites {*i*_1_, *i*_2_, …, *i*_*k*_}, a relation corresponds to a set *M*_*k*_[*i*_1_, …, *i*_*k*_] consisting of k-tuples (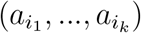), representing all the constellations that satisfy the “hidden” relation. We shall refer to 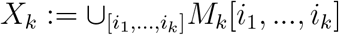, as the set of *k*-motifs or *motifs*. Constellations are projective: 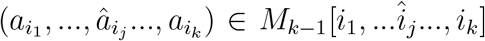 for any *j* ∈ {1, …, *k*}, where 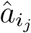 expresses the fact that 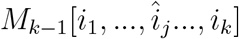 is omitted, i.e. any (*k −* 1)- tuple induced by an *M*_*k*_[*i*_1_, …, *i*_*k*_] element, is a corresponding *M*_*k−*1_[*i*_1_, …, î_*j*_…, *i*_*k*_]- tuple. The projectivity reflects the fact that, by construction, any sub-motif will be observed as an induced co-evolutionary dependency.

Suppose the set of all motifs, *X* := ∪_*k*_X_*k*_, is given. Its simplices encapsulate relations that are represented by mutational constellations within the MSA and it is natural to endow them with weights, representing the number of distinct constellations realizing them. Accordingly, *X* gives rise to a weighted simplicial complex [5] over the set of columns defined as follows: [*i*_0_, …, *i*_*k*_] *∈ X* is a *k*-simplex of weight *w* if and only if |*M*_*k*_[*i*_1_, …, *i*_*k*_]| = *w >* 0.

We now adopt the following perspective: suppose we are given an MSA providing consistent labelings of the sites relative to a reference sequence and suppose a family of motifs (*M*_*k*_[*i*_1_, …, *i*_*k*_])_*k*_ exists, but is not known to the observer. Then the MSA allows to obtain information about maximal motifs, representing the sets of co-evolving sites. Depending on size and composition of the MSA, as well as errors introduced by constructing the MSA the “true” motif complex, i.e. the collection of all blocks having implicit connection, can only be approximated.

To this end we construct simplices starting from vertices (sites) (0-simplices) to maximal simplices. These maximal simplices are of central relevance, since they represent maximal collections of co-evolving positions, which include^1^ the crucial functional units in the virus genome.

In view of 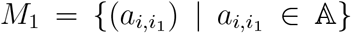 there exist, no *a priori* constraints on the selection of the sites of the motif complex. We shall select sites within the MSA that play a distinguished role in the *evolutionary dynamics* of the sequence sample:

- sites contained in competing variants within the multi-sequence alignment or
- sites exhibiting significant variation for intrinsic, biochemical reasons.

In order to recover the motif complex, we employ measures of nucleotide diversity and co-evolution distances as follows: first we identify the critical sites where selection induces evolutionary variation and secondly we quantify those pairs of sites that co-evolve.

First, we use Hamming distance and explicitly incorporate a particular class of relations that is induced by permutations and secondly we employ entropy and mutual information. We remark, that in the latter case, although not explicitly encoded, permutation induced relations again emerge.

## The motif complex of SARS-CoV-2

In this section we consider the motif complex of SARS-CoV-2. Using results from section *Materials and Methods* we approximate the complex and discuss alerts, actual clusters and true and false positives.

By construction, the motif complex does not allow us to draw conclusions as for which motifs will constitute a “problem”; this can only be achieved by detailed biological analysis. Short of providing such an analysis, the identification of motifs is critical and of timely value because of

- a dramatic reduction of the number of potentially relevant sites from the order of 10^4^ to 10^2^,
- rapid detection of collections of sites that constitute potential threats.

We next specify *alerts*. To this end, we refer to a site as newly emerging ((+)-site), if it changed its activity state from being inactive to active within the MSA and (*−*)- site, otherwise. Here, the activity is measured by the average number of pairwise different nucleotides at a specific site, denoted by *D*(*i*). We approximate motifs as detailed in Section *Materials and Methods*, referring to these as clusters. Given a discrete time series a particular cluster, *M* splits 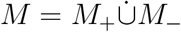.

**Alert**: *is the emergence of a motif, M, such that* |*M* | *≥* 5 *and ρ*_*M*_ = |*M*_+_|*/*|*M* | *≥* 0.5. A cluster triggering an alert is referred to as predicted positive.

A cluster of size at least five containing at least one (+)-site is referred to as *actual*. Actual clusters provide the background for the ROC-curve provided in Subsection *Alerts: genericity and ROC-curves*. As we shall see, alerts are not random and, as the below case studies show, only few are triggered at a given time, involving on the order of 10 positions.

An alert, i.e. a predicted positive can be either a true or false positive. This is decided via the following criterion: in case more than 70% of the sites contained in the cluster, irrespective of them being (+) or (*−*)-sites, are later^2^ confirmed to be characteristic mutations of a single VOI/VOC, we consider an alert a true positive and a false positive, otherwise. We give a detailed analysis of the dependency of the alert criterion on its key parameter *ρ*_*M*_. The alert criterion, specified above, is arguably *ad hoc*. It turns out that while it can be optimized, the optimization does not increase true positives, but decreases false positives, see Subsection *Alerts: genericity and ROC-curves for details*.

Finally, we draw the attention to an additional feature of motifs: when mapped onto specific VOCs and VOIs, motifs provide deeper insight in how characteristic mutations organize, which in itself aids the biological analysis.

In Tab. 1 we compare the time of detection of critical motifs, corresponding to VOC/VOIs to the time of (a) WHO designation [48] as being of concern/interest and (b) Pango lineage designation [39].

**Table 1.**
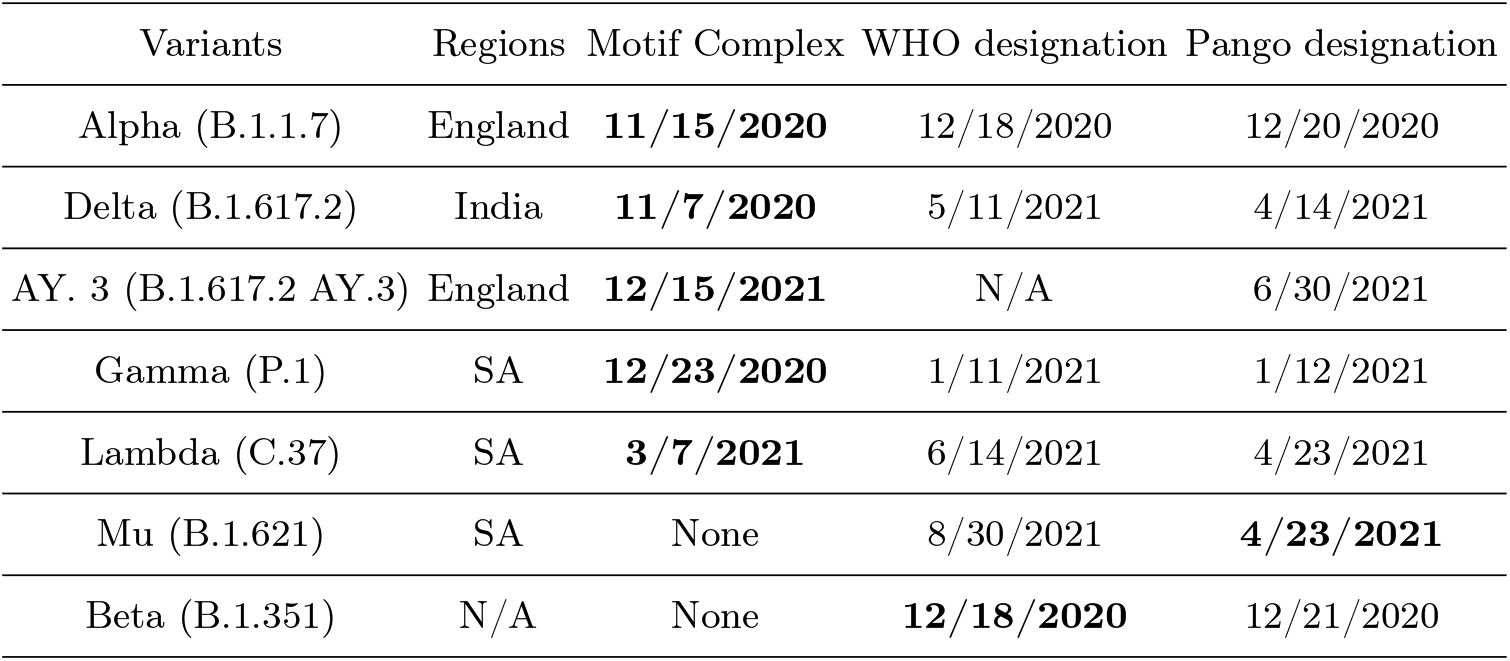
Rapid detection: motif-based alert, WHO and Pango designation. Motif detection: the date an alert is observed. WHO designation: the date the respective variant was declared to be of concern/interest. Pango designation: the date a lineage was assigned to the respective variant according to Pango designation website.

| Variants | Regions | Motif Complex | WHO designation | Pango designation |
| --- | --- | --- | --- | --- |
| Alpha (B.1.1.7) | England | **11/15/2020** | 12/18/2020 | 12/20/2020 |
| Delta (B.1.617.2) | India | **11/7/2020** | 5/11/2021 | 4/14/2021 |
| AY. 3 (B.1.617.2 AY.3) | England | **12/15/2021** | N/A | 6/30/2021 |
| Gamma (P.1) | SA | **12/23/2020** | 1/11/2021 | 1/12/2021 |
| Lambda (C.37) | SA | **3/7/2021** | 6/14/2021 | 4/23/2021 |
| Mu (B.1.621) | SA | None | 8/30/2021 | **4/23/2021** |
| Beta (B.1.351) | N/A | None | **12/18/2020** | 12/21/2020 |

We next discuss alerts and how their underlying motifs relate to VOCs/VOIs in terms of four case studies: the Alpha (England), Delta (India), Delta AY.3 sub-lineage (US), and Mu variant in SA. We shall approximate motifs as detailed in Section *Materials and Methods*, restricting ourselves to an analysis based on *P* - distance combined with *k*-means clustering. Approximations involving *J* distance and HCS-clustering are presented in the Supplemental Materials. We will show that we can rapidly identify clusters of co-evolving active sites–only later to be recognized as characteristic mutations of a respective variant, see Tab. 1.

### Alpha and AY.3 (England)

Following the protocol described in Section *Materials and Methods*, we analyze SARS-CoV-2 genomic data collected in England. We display alerts on a weekly basis ranging from week 1, November, 2020, to week 4, August, 2021, see Fig. 1. We observe six alerts, five of which *a posteriori* are confirmed to mark the emergence of a VOI/VOC.

**Figure 1.**
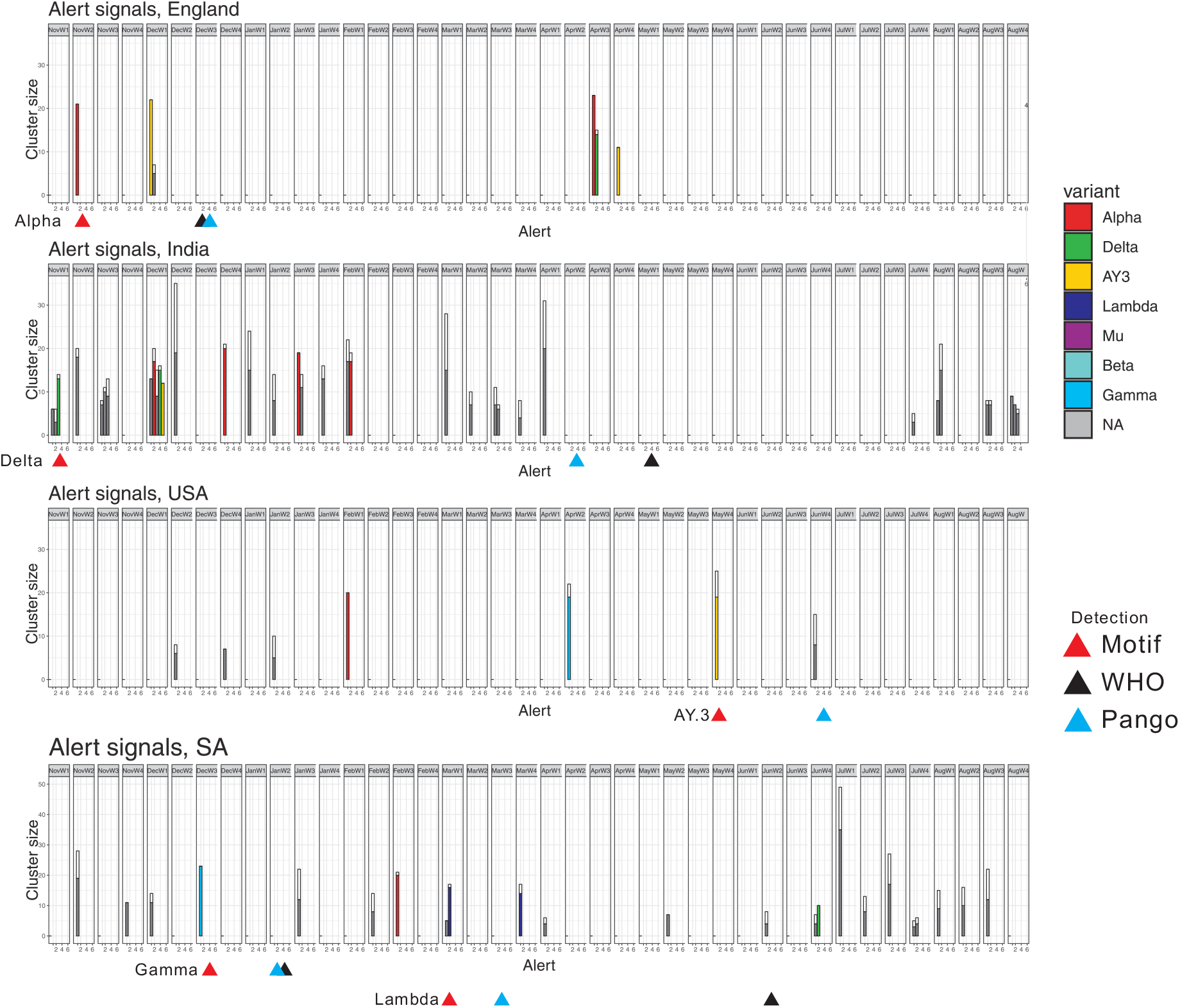
Motif-induced alerts: each vertical represents a cluster satisfying our alert criterion, where the height denotes the size of the cluster and the color portion of a vertical bar representing the ratio of (+)-positions within the cluster. An alert mapping onto a particular VOI/VOC is colored accordingly and colored grey, otherwise. Triangles label the first week of detection of the motif-based alert, WHO and Pango designation, see Tab. 1.

Week 2, November 2020, a single cluster composed by 21 (+)-positions emerges, see Fig. 1. These sites are later confirmed to be among the 28 characteristic mutations of the Alpha variant. The motif complex organizes these 28 mutations into three groups, see Fig. 2: the mutations C241T, C3037T, C14408T, and A23403G (*C*_1_, gray), the mutations G28881A, G28882A, G28883T (*C*_2_, blue), and the remaining 21 mutations (*C*_3_, red).

**Figure 2.**
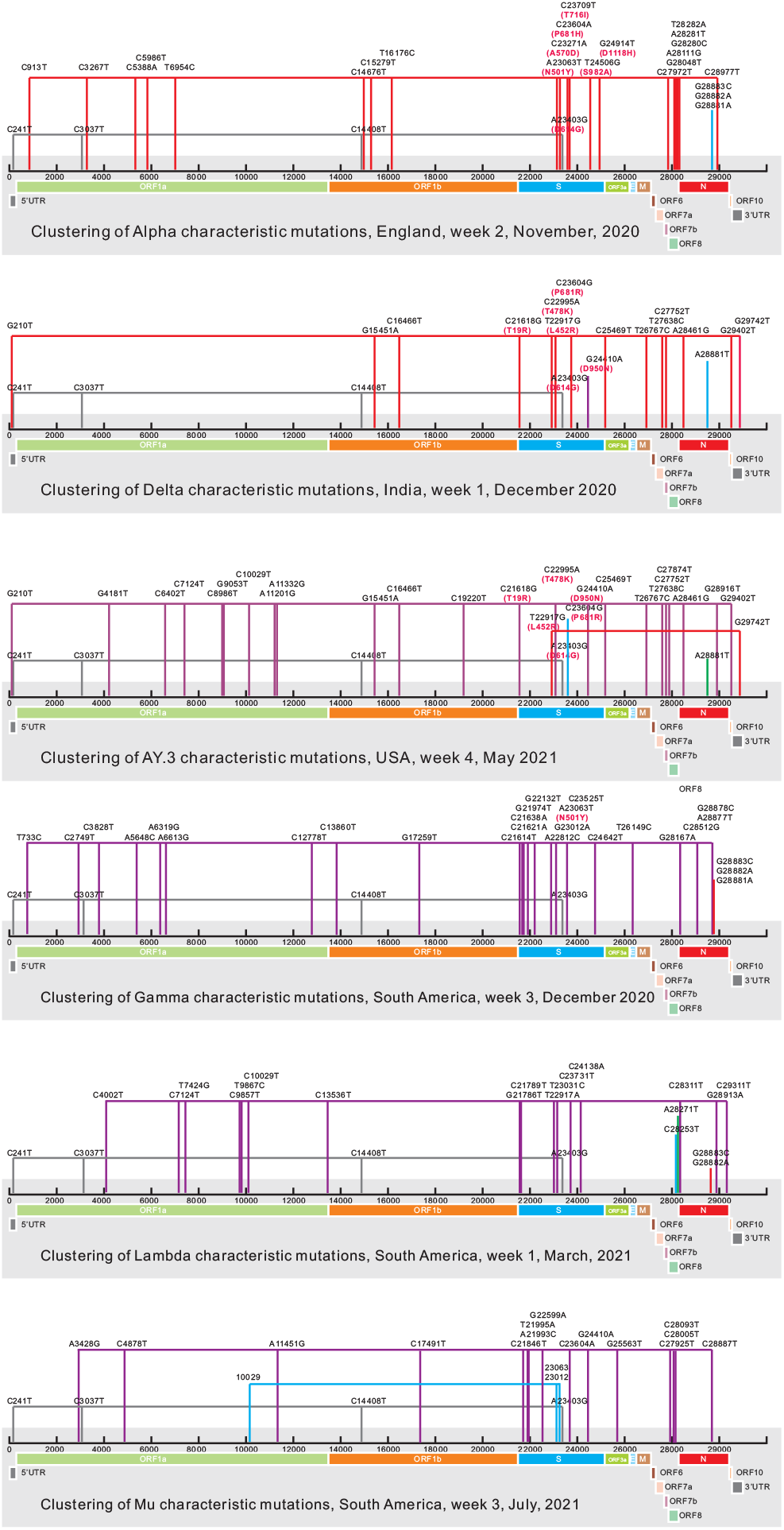
Motif-induced blocks of Alpha, Delta, AY.3, Gamma, Lambda, and Mu variants: associated sites and characteristic mutations on the viral genome. Sites connected on the same level correspond to clusters/motifs.

The four *C*_1_-mutations, responsible for the D614G amino acid change on the spike protein, first emerge simultaneously in February 2020 and dominate the viral population by June 2020. *C*_1_ is found in all variants after June 2020 and our approach always clusters these. *C*_2_ is present in the viral population since February 2020, but does not become dominant. Its three positions correspond to two adjacent codons and the mutations cause various amino acid changes R203K and G204R on the N protein, also found in other variants, such as Gamma, and our approach correctly clusters them. *C*_3_ is first observed in the 2nd week of November. Its positions are exclusively co-evolving in Alpha.

Week 1, December, see Fig. 1 England, an entire cluster containing 22 (+)-sites emerges, later confirmed as the characteristic mutations of the Delta sub-lineage AY.3, 11 of which being shared with Delta and are in fact characteristic mutations of the Delta variant.

In summary, an alert marks the rise of the Alpha variant in a timely fashion, at least four weeks earlier that the WHO and Pango designations. In addition, motifs reveal insight into the organization of characteristic mutations into co-evolving blocks: the 28 defining mutations of Alpha are seen to split into three blocks, see Fig. 2.

### Delta (India)

We observe 35 alerts from November 2020 through August 2021 in India, 7 of which are confirmed as the emergence of a VOI/VOC. Specifically, we have three alerts in week 1 of November, 2020, see Fig. 1 India. One of these corresponds to a cluster of 14 sites, 13 of which being (+)-sites and later confirmed to be among the 20 characteristic mutations of the Delta variant. These 13 positions become inactive in the following three weeks.

Five alerts are triggered in week 1, December: (a) an entire newly emergning cluster of size 13, not mapped to any VOI/VOC, (b) a cluster of 20 sites, 17 of which are (+)-sites mapping to the Alpha variant, (c) a cluster of 15 sites, 9 of which are (+)-positions, (d) a cluster of 16 sites 15 of which are (+)-positions mapping to the Delta variant, and (e) an entire newly emerging cluster of 12 (+)-sites, mapping to the Delta AY.3 sub-lineage.

Delta comprises 20 characteristic mutations, see Fig. 2. Motifs organize these into four blocks representing the co-evolutionary footprint of Delta, week 1, December: C241T, C3037T, C14408T, and A23403G (*C*_1_, gray), C23604G (*C*_2_, blue), G24410A (*C*_3_, purple) and the remaining 13 mutations (*C*_4_, red), see Fig. 2. The *C*_1_-mutations have been discussed in the previous section. C23604G (*C*_2_) is contained in the Alpha variant and emerged earlier. The site corresponding to G24410A (*C*_3_) emerged in a different cluster in the 1st week of December, 2020, similar to the *C*_4_-sites, that appear in a cluster at the same time.

The characteristic mutations of the Delta variant do not emerge at once, they arise over the period from November 2020 through April 2021, see Fig. 3. The motif complex provides deeper insight into the co-evolutionary footprint during their development.

**Figure 3.**
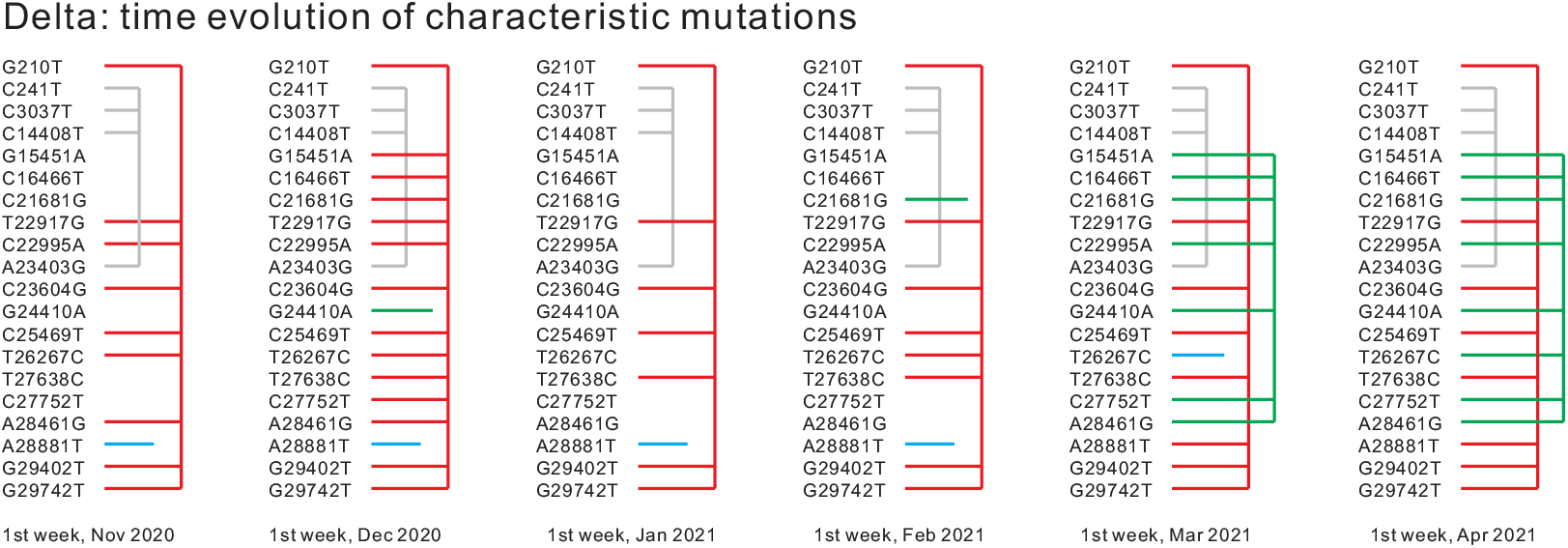
The evolution of the co-evolutionary footprint of Delta: India, from November 2020 to April, 2021, based on *P*-distance and *k*-means clustering. Sites not clustered are inactive at the respective time (*D*(*i*) *≤* 0.1, see Materials and Methods).

Delta exhibits 20 characteristic mutations, including the *C*_1_-mutations (grey) also found in any current variant. We shall consider here the organization of the remaining 16 mutations. In November, we find that 10 mutations among the 16 characteristic mutations are active, and 9 cluster, leaving A28881T isolated. A28881T is a *C*_2_-mutation which emerged earlier and can be found in other variants. In December, all 16 characteristic mutations are active, 14 forming a cluster, while A28881T and G24410A are isolated. G24410A is an essential mutation on the spike protein region, resulting in D950N amino acid substitution. The situation is somewhat similar during November, January and February, in March, however, all 16 characteristic mutations become active and form three clusters: of size 8 (red), of size 7, composed of newly emerging mutations (green), and of size one, T26267C (blue). In April, the situation is similar to March. Two large clusters are observed, where T26267C merges into one of them (green). We conduct the same type of analysis via the HCS-method on the April data. Here the 16 characteristic mutations partition into 8 clusters: two large clusters of size 5 and 6 small clusters of size 1. Accordingly, both *k*-means and HCS clustering split the characteristic mutations of Delta into two large clusters.

### AY.3 (USA)

AY.3 is a sub-lineage of Delta and the characteristic mutations of Delta form a subset of its characteristic mutations. AY.3 exhibits 31 characteristic mutations, 11 of which belong exclusively to the AY.3 sub-lineage, see Fig. 2. In the USA we observe 7 alerts between November 2020 through August 2021, 3 of which were confirmed to mark the emergence of a VOI/VOC.

Week 1, February, an entire cluster composed of 20 (+)-sites, emerges, covering, with the exception of C23604A, the *C*_3_ mutations, see Fig. 2. Week 2, April, a cluster of size 22, 19 of which being (+)-sites emerges and all mapping to the Gamma variant. Week 4, May, a cluster of 25 sites emerges, 19 of which being (+) and are later confirmed to be among the 31 characteristic mutations of the Delta AY.3 sub-lineage. Within this cluster, 10 sites correspond to characteristic mutations exclusive to the Delta AY.3 sub-lineage, the only exception being C10029T, which emerges earlier in the US.

### Lambda and Mu (SA)

From November, 2020 through August, 2021, we display the alerts in SA, see in Fig, 1. We observe 23 alerts, 5 of which being confirmed as the emergence of a VOI/VOC.

Week 3, December, an entire cluster containing 23 (+)-sites emerges, mapping to the Gamma variant. Week 3, February, a cluster containing 22 sites 21 of which being (+)-sites and mapping to the Alpha variant. Week 1, March 2021, a cluster of size 17 emerges, containing 16 (+)-positions, later confirmed as to constitute the defining mutations of the Lambda variant, see Fig. 2. Week 4, March, a cluster of size 17 emerges, containing 14 (+)-sites and mapping to the Lambda variant. Week 4, June, an entire cluster composed of 10 (+)-sites emerges, mapping to the Delta variant.

The Lambda variant is detected at an early stage, although in March 2021 the prevalence within the viral population of Lambda is low (*<* 5%) and not yet assigned as a lineage by Pango, we identify a distinguished co-evolutionary signal mapping to the Lambda variant 6 weeks “ahead of time”.

The co-evolutionary footprint of the Lambda and Mu mutations, see Fig. 2 show that within Lambda there is a distinguished cluster of size 16, while the remaining three clusters are of size 2, 1, and 1 respectively. Mutations corresponding to the positions in the cluster of size 2, G28882A and G28883C, are also found in Alpha and Gamma, while the one corresponding to the position in the cluster of size 1, A28271T, is found in the Delta AY.3 sub-lineage. The remaining cluster of size 1 is C28253T. The Mu variant exhibits similar feature, a distinguished cluster of size 15 and two smaller clusters are of sizes 3 and 4 respectively.

Our approach does *not* issue an alert of the emergence of the Mu variant. This is by construction: the characteristic mutations of the Mu variant are a recombination of mutations present in other variants such as Alpha, Gamma and Delta. As such these mutational sites are already active in the population, and therefore they are not considered as (+)-positions.

### Motif reconstruction

In this section we present a comparative analysis of the four motif reconstruction methods, obtained by combining *P* - and *J*-distance with *k*-means and HCS-clustering. The following analysis is based on the SARS-CoV-2, whole genome data, for July, USA, collected from GISAID.

When constructing the similarity graph, which is the input for the HCS-clustering algorithm, we set the threshold for column selection homogeneously to be *D*(*i*) *>* 0.1, where *D*(*i*) denotes the diversity of site *i*, see Materials and Methods. The parameters for *P*- and *J*-distance are set 0.1 and 0.45 respectively, i.e. an edge is placed in the similarity graph for any pair of columns having *P*-distance *<* 0.1 and *J*-distance *<* 0.45, respectively.

We present the result in form of an *m × m* matrix, *T* = (*t*_*i,k*_) where *m* denotes the number of sites such that *D*(*i*) *>* 0.1, see Fig. 4. An entry *t*_*i,k*_ is drawn black if (*i, j*) are both contained in a distinguished cluster and white, otherwise. Since distances are symmetric, we can present data regarding two distances with respect to a particular clustering method within a single matrix, bi-partitioning it into its upper-left and lower-right triangular blocks.

**Figure 4.**
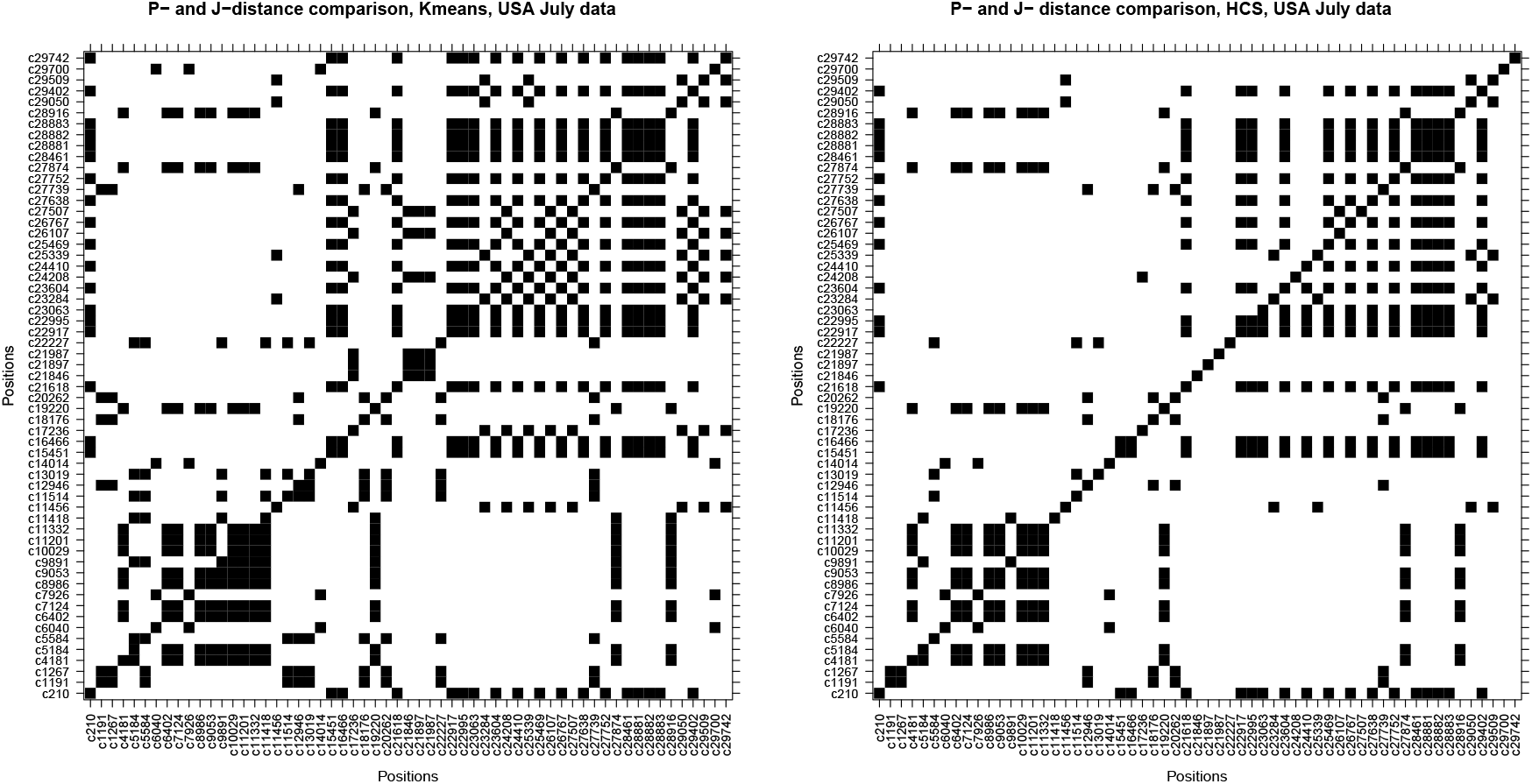
*k*-means and HCS clustering based on *J* - and *P*-distance: the *x*- and *y*-axis display active columns (*D*(*i*) *>* 0.1). If the *i*th and *j*th positions belong to the same cluster, (*i, j*) is colored black and white, otherwise. LHS: *J*-distance and *k*-means clustering (upper triangle), *P*-distance and *k*-means clustering (lower triangle). RHS: *J*-distance and HCS clustering (upper triangle), *P*-distance and HCS clustering (lower triangle).

**Figure 5.**
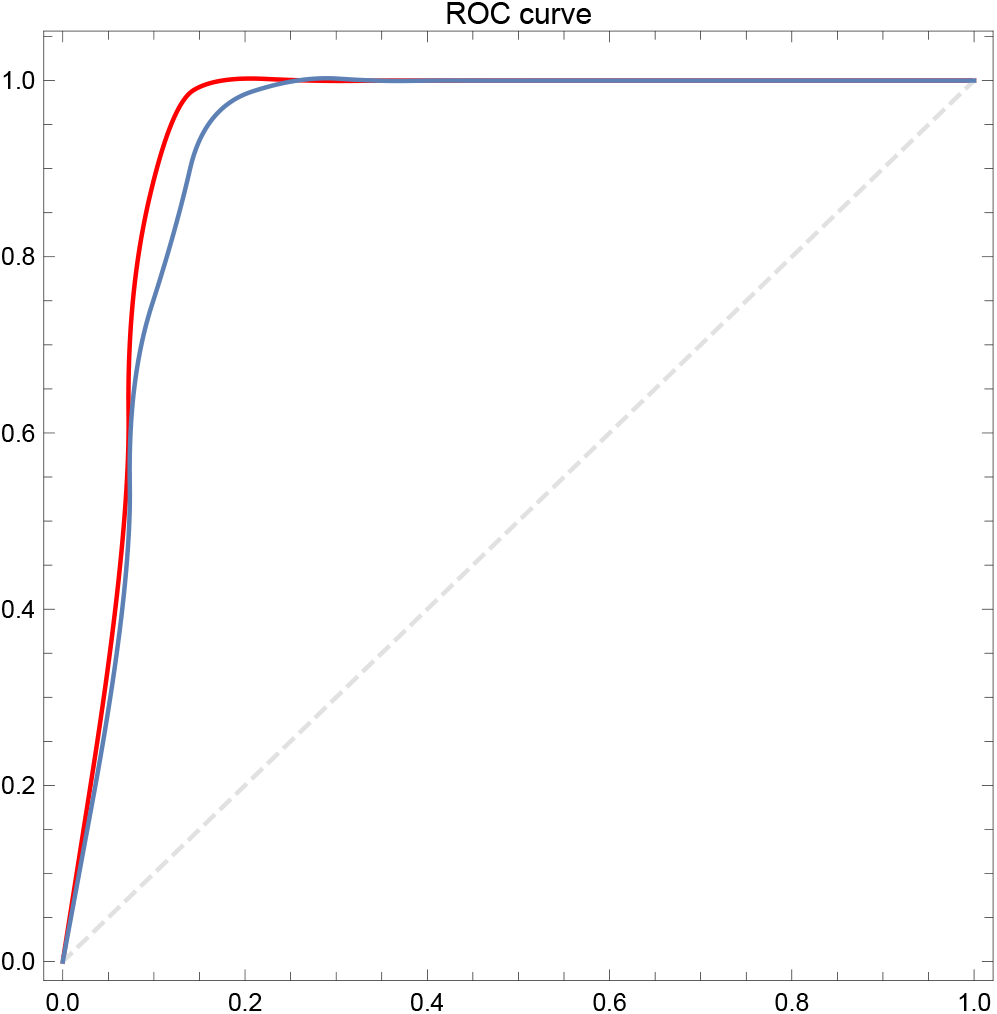
ROC curves of alerts: *x*- and *y*-axis denote the false and true positive rates *FPR* and *TPR*. Each point corresponds to a specific *θ*, i.e. the fraction of newly emerging sites within the cluster triggering the alert, ranging from 1 to 0 (left to right). We display the ROC-curves in case of true positives corresponding to 70% (red) and 50% (blue) of the alert-cluster representing characteristic mutations of a VOC.

The matrices exhibit a high degree of symmetry, indicating that *k*-means clustering and HCS-clustering produce similar results for *P* - and *J*-distance. This suggests that both metrics capture co-evolution in the SARS-CoV-2 data.

### Alerts: genericity and ROC-curves

We first analyze the dependency of the alert protocol on metric and clustering method. To this end we compare all combinations of *P* - and *J*-distances with *k*-means and HCS-clustering. The following analysis is based on GISAID SARS-CoV-2 whole genome data, collected from November 2020 through August 2021, for England, USA, India and SA.

For all four cases, we set the threshold for activity to be *D*(*i*) *>* 0.1. For construction of the similarity graph, the input for the HCS-clustering algorithm, the threshold parameters are set to be 0.1 and 0.45, for *P* - and *J*-distance, respectively. All four combinations produce similar signals, see Fig. S1–S3 in the SM. Specifically, they all induce alerts that mark the emergence of the Alpha and the AY.3 variants in England. Furthermore, the emergence of the Gamma variant is also captured by all variations. This suggests that the co-evolutionary signals in the SARS-CoV-2 data can be derived by a variety of methods.

However, we also observe systematic differences: in general, HCS-clustering tends to produce smaller clusters when compared to the *k*-means method. This is due to the fact that forming a highly connected component is a restrictive condition. We also observe that P-distance produces slightly more signals compared to J-distance.

To study the diagnostic capability of alerts, we perform a receiver operating characteristic (ROC) analysis [13]. The key parameter here is the threshold of (+)- mutations, *θ*. In case the fraction of (+)-mutations contained in the cluster exceeds *θ*, an alert is triggered and the corresponding cluster is considered a predicted positive. By construction, the total number of predicted positive clusters is a monotoneously decreasing function of *θ*.

Any predicted positive cluster corresponding to a VOC/VOI, is considered a true positive and a false positive, otherwise. Let *TP* and *FP* denote the total number of true positive and false positive clusters, respectively. Then the true and false positive rates *TPR* and *FPR* are given by *TP/P* and *FP/N*, where *P* and *N* denote the numbers of actual positives and negatives, respectively, where an actual cluster consists of at least 5 sites and contains at least one (+)-site. The latter guarantees that the cluster is only counted once. When at least 70% (50%) of an actual cluster correspond to the characteristic mutations of a certain VOC/VOI, this cluster is considered to be associated with that VOC/VOI, and contributes to *P*. If an actual cluster does not correspond to any VOC/VOI, then it contributes to *N*.

We note that the ROC curve depends on (*P, N*) being correctly recognized. If, for instance, an only later to be declared VOC is not taken into consideration all fractions will change. Each *θ* induces a tuple (*FPR, TPR*). Varying *θ*, produces the ROC curve [13], where the *x*-axis and *y*-axis represent the *FPR* and *TPR*, respectively.

Integrating over all data, that is, considering all geographical locations, we observe a total of 163 actual clusters (each consisting of at least 5 sites and containing at least one (+)-site). Among them, *P* = 20 correspond^3^ to a VOC/VOI and are accordingly considered to be actual positives. The remaining *N* = 143 do not correspond to a VOC/VOI and are considered to be actual negatives. We point out that none of the actual clusters is associated with Mu. In fact, the characteristic mutations of Mu distribute over multiple actual clusters, Mu-sites are to large extend a recombination of sites of other variants. The ROC curve displays the trade-off between sensitivity (*TPR*) and specificity (1 *− FPR*): a random classifier is expected to produce points lying close to the main diagonal.

## DISCUSSION

Co-evolution represents a crucial footprint of evolution. Genomic adaption is to large extent not facilitated via isolated but collections of mutations. Functionally connected mutations changing simultaneously is a key indicator of relevant viral dynamics contained in the MSA. Co-evolving mutations are indicative of the existence of noteworthy biological mechanisms inducing differentials in the evolutionary dynamics. However, they do not allow us to infer details about such biological mechanisms.

The motif complex introduced here detects maximal sets of sites that experience selection pressure –and this is the crucial point– *as a collective*. This amounts to identify differential changes within the MSA, providing information about the viral “heartbeat”. We have shown that this pressure leads to a small number of mutational constellations that appear as distinguished patterns within the MSA. Thus the method represents a significant reduction in data and facilitates subsequent biological analysis.

In contrast to the current approach, the motif-complex does not require any *a priori* assumptions as it is a bottom-up approach. In contrast, the current approach of defining a lineage or variant is based on its location in the phylogenetic tree. Determination of branch points can be biased and there is no clear boundary between lineages since their characteristic mutations can overlap. Motifs provide a new way of partitioning mutations. A position can only be clustered in a group at a time. A cluster is possibly representing a functional block, and the current defining variant is a combination of these functional blocks. For example, all variants contain the cluster of mutations C241T, C3037T, C14408T, and A23604G.

The more sequences the MSA contains the more easily such constellations are observed. Quality and quantity of sequence data affect the fidelity of approximation. Sparse data, on the other hand, as it is the case for India, have a detrimental effect, since the MSA does not provide a sufficient basis for the reconstruction of the “true” motifs. England and USA surveillance have a higher number of sequences [4], allowing for the reconstruction of the motif complex with much higher fidelity. We observe that retrospective alerts based on England and USA GISAID-data appear rarely and typically correspond to VOCs/VOIs. On the contrary, Columbia, where the Mu variant is emerging from, has much fewer sequences due to global disparities in sequence surveillance [4]. It is worth mentioning that sequence surveillance has an impact on the motif complex detecting the Mu variant in Columbia.

Alerts are based on the approximation of motifs. Irrespective of the particular parameterization, observing a motif is non-random since it is produced by selection pressure acting on a collective of sites. Evolution in absence of selection, i.e. on a flat landscape produces lineages and clusters^4^ [9] but never exhibits *P* - or *J* - distances small enough to form even a motif composed by only two positions. Thus, in contrast to lineages, the existence of motifs is tantamount to the existence of selection pressure.

A set of co-evolving sites can originate from a variety of biological scenarios. Particularly relevant events closely connected to motifs are for instance functional blocks. These induce subsets of motifs since the method cannot rule out that only a core of sites is directly relevant for the underlying functionality, while remaining sites are “carried along” by founder effect or other mechanisms. It is, for instance, easily conceivable to have two functional units forming a motif, where the existence of one excludes the other. In any case, motifs represent a dramatic data-reduction, since there are only a few of them and they consist typically of *≤* 20 sites.

Even if all motifs would correspond to functional blocks, not all would result in VOCs or VOIs. The emergence of these depends on the viral dynamics itself, strain competition, as well as external factors such as selection pressures exerted via vaccinations or social distancing.

The approximation of the motif-complex of an MSA identifies the co-evolutionary relations between sites on the genome. This constitutes key data, that can be utilized to get an instant read on the evolutionary dynamics within the viral sample. Our results show that this information is instrumental for the early detection of differential changes in the dynamics of the sequences in the MSA. Such signals can be detected before the adaption of a variant is complete. As the Delta variant exemplifies, this adaption can be a months-long process, through which sites configure themselves into an optimal constellation in multiple steps, each of which leaving its co-evolutionary footprint.

Combining the maximal simplices or clusters with a phylogenetic analysis provides deeper insight into how VOCs are organized, see Fig. 2. As a result we can show that the characteristic mutations representing a VOC split into distinguished components of co-evolving sites.

The concept of alerts works well to achieve the goals of the method. Over a wide parameter range we produce a true positive rate of 1, i.e. no VOI/VOC is missed, which motivates the title of this contribution. However, that is not to say that the method is not without its limitations. This has to do with our notion of “universe”, i.e. the set of actual clusters and what amounts to a positive. The Mu variant does not appear to cover a sufficient fraction of any actual cluster and is therefore not counted as an actual positive. Consequently, it is fair to say our method genuinely hits all VOI/VOC that exhibit a significant fraction of *de novo* mutations. Mu can be considered as a recombination of sites, that are present in other variants. Mu-sites are thus distributed over a large number of actual clusters, which leads by construction to its exclusion from the set of positives. Similarly our method does not detect signals mapping to the Beta variant in the four countries/regions, see Tab. 1.

The framework generates false positives at a rate below 0.2. This is acceptable in view of the fact that the underlying clusters are small and only emerge within a week. We remark that our measure of false positives includes clusters that may under different circumstances not be false positives at all. These clusters do represent a critical threat, but for reasons of strain competition, founder effect, or external measures such as social distancing, this threat never materializes.

## Materials and Methods

### First approximation

We first compute the nucleotide diversity of a column, *i*, i.e its average Hamming distance [19] 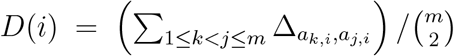, where Δ_*i,k*_ = 1 *− δ*_*i,k*_ and *δ*_*i,k*_ is the Kronecker symbol.

We proceed by approximating its 1-simplices. To this end we consider all permutations *τ* : 𝔸 *→*𝔸 and make the Ansatz, see Fig. 6:

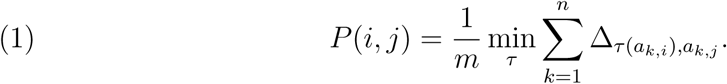

**Figure 6.**
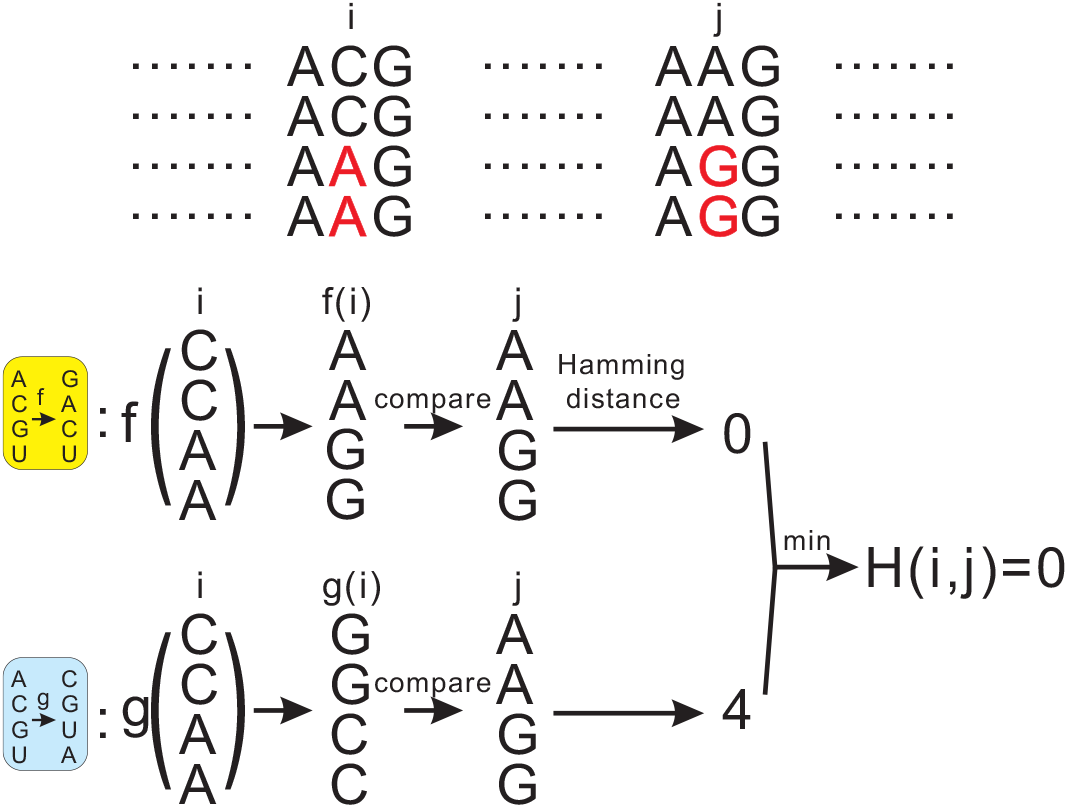
Permutation-induced relations: at site *i* and *j* suppose *v*_*i*_ = CCAA and *v*_*j*_ = AAGG, respectively. Let *g* map e to e, A to C, C to G, G to U, and U to A. Then *g*(*v*_*i*_) = GGCC and the Hamming distance between *g*(*v*_*i*_) and *v*_*j*_ is four. For *f* mapping e to e, A to G, C to A, G to C and U to U, the Hamming distance between *f* (*v*_*i*_) and *v*_*j*_ is zero and *P* (*i, j*) = 0, accordingly.

In the context of a noisy data-set, *P* (*i, j*) can be viewed as to reverse-engineer the dependencies induced by permutations between the two columns. Such permutations produced a restricted, yet relevant collection of binary relations. For instance, relations like identity or complementarity can readily be expressed via such mappings. *P* (*i, j*) satisfies by construction *P* (*i, j*) = *P* (*j, i*) and the triangle inequality *P* (*i, h*) *≤ P* (*i, j*) + *P* (*j, h*). That is *P* (*i, j*) is a pseudo-metric and since there are 5! permutations, *P* (*i, j*) can be computed easily. *P* (*i, j*) is completely determined by the joint distribution *p*_*i,j*_(*x, y*) of pairs of nucleotides, namely *P* (*i, j*) = 1 *−* max_*τ*_*Σ* _*x*_ *p*_*i,j*_(*x, τ* (*x*)).

We are now in position to approximate the motif complex based on *D*(*i*) and *P* (*i, j*) as follows: first, we define the 0-simplices to be the columns *i* such that *D*(*i*) is greater than the threshold *h*_0_, *D*(*i*) *> h*_0_. Second, we define the 1-simplices as follows: a pair of columns (*i, j*) is a 1-simplex if *P* (*i, j*) is smaller than the threshold *E, P* (*i, j*) *< E*. In view of the property if *P* (*i, j*) = *P* (*j, h*) = 0, then *P* (*i, h*) = 0, we shall thirdly approximate the higher dimensional *k*-simplices for *k ≥* 2 as follows: any *k* + 1 columns [*i*_0_, *i*_1_, …, *i*_*k*_] form a *k*-simplex of the motif complex if any pair of these columns forms a 1-simplex. These *k*-simplices for *k ≥* 2 can be approximated by means of cluster analysis or alternatively, extractions of highly connected subgraphs of a similarity graph induced by *P* (*i, j*) via the HCS-algorithm.

### Second approximation

We determine the 0-simplices of the complex via the *Shannon entropy, H*(*i*), of a site *i*, given by [43] *H*(*i*) = *− Σ* _*x∈*𝔸_ *p*_*i*_(*x*) log_2_ *p*_*i*_(*x*), where the units of *H* are bits, and *p*_*i*_(*x*) is the probability of the nucleotide *x* appearing in column *i*. The entropy *H*(*i*) has been widely utilized to quantify the diversity of nucleotides at position *i* in a population of sequences [43, 7]. In case of entropy, we shall construct a distance via *joint entropy* and *mutual information* as follows: the *joint entropy H*(*i, j*) of two sites *i* and *j* is defined as

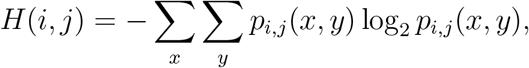

where *p*_*i,j*_ denote the joint distribution of columns *i* and *j*, i.e., *p*_*i,j*_(*x, y*) specifies the probability of pairs of nucleotides (*x, y*) *∈* 𝔸 *×* 𝔸. Clearly, the marginal probability distributions for columns *i* and *j* are given by *p*_*i*_(*x*) = Σ _*y*_ *p*_*i,j*_(*x, y*) and *p*_*j*_(*y*) = Σ_*x*_ *p*_*i,j*_(*x, y*), respectively.

The *mutual information I*(*i*; *j*) between sites *i* and *j* is the relative entropy between the joint distribution *p*_*i,j*_(*x, y*) and the product distribution *p*_*i*_(*x*)*p*_*j*_(*y*):

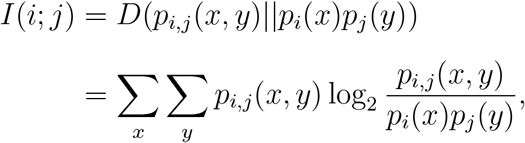

where *D*(*p*||*q*) denote the *relative entropy* or *Kullback-Leibler divergence* from the distribution *p* to the distribution *q* [28]. *The mutual information I*(*i*; *j*) quantifies the amount of information shared by two columns *i* and *j*.

Then the *J-distance J* (*i, j*) between two sites *i* and *j* is given by

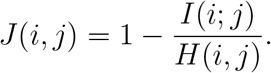

The J-distance represents the information-theoretic counterpart of the Jaccard distance [22, 7]. *J* (*i, j*) satisfies the following properties: 0 *≤ J* (*i, j*) *≤* 1, *J* (*i, j*) is a pseudo metric, i.e. *J* (*i, i*) = 0, *J* (*i, j*) = *J* (*j, i*) and the triangle inequality *J* (*i, j*) *≤ J* (*i, k*) + *J* (*k, j*). Furthermore, *J* (*i, j*) is *scale invariant*, that is *J* (*i, j*) is independent of the population size, and uniquely determined by the joint distribution of pairs of bases.

We are now in position to approximate the motif complex based on the information-theoretic measurements of the columns. First, we define the 0-simplices to be the columns *i* such that *H*(*i*) *> h*_0_ and a pair of columns [*i, j*] is a 1-simplex if *J* (*i, j*) *< γ*. In view of *J* (*i, j*) = 0 and *J* (*j, k*) = 0, implying *J* (*i, k*) = 0, for *k ≥* 2 any *k* + 1 columns form a *k*-simplex [*i*_0_, *i*_1_, …, *i*_*k*_] if any pair of these columns forms a 1- simplex.

It is worth mentioning that *P* - and *J*-distance are both closely tied to permutations generating the motifs. While this holds obviously for *P* (*i, j*), it follows for *J* (*i, j*) from the fact that if *J* (*i, j*) = 0, then there exists a bijection *τ* on the alphabet such that, for each *x, p*_*i,j*_(*x, τ* (*x*)) = *p*_*i*_(*x*) = *p*_*j*_(*τ* (*x*)) and *p*_*i,j*_(*x, y*) = 0, otherwise.

### Cluster analysis

For the clustering analysis, we employ *k*-means clustering [30] and the highly connected subgraphs (HCS) clustering [21]. While the optimal cluster number via *k*-means clustering is determined via the within sum of squares distribution, the similarity graph via HCS is constructed via a certain threshold on the co-evolution distances. Comparing the two methods, we note that there is a trade-off: while *k*-means clustering requires the number of clusters to be specified, but no other parameters, HCS-clustering requires a threshold to be set for the construction of the similarity graph, while being indeterminate on the number of clusters.

Either method produces an approximation of the motif complex, whose high dimensional simplices correspond to clusters of co-evolving sites (by means of a variety of mutational constellations). The resulting clusters of co-evolving sites are of crucial importance for detecting blocks that are experiencing selective pressure, representing in some sense the evolutionary pulse within the multiple sequence alignment. *k*-means clustering [30] is an unsupervised machine learning algorithm of vector quantization, aiming to partition *n* observations into *k* clusters in which each observation belongs to the cluster with the nearest mean to the cluster center. The problem is in general NP-hard [2], but efficient heuristic algorithms are available and can achieve a local optimum [20, 42]. As the number of clusters *k* is part of the input, the first step is to determine a suitable *k*. Here we use the gap statistic method [45], with maximum *k* = 30, to determine the optimum *k*. The analysis is performed by the factoextra package in R [29, 23].

We also perform the highly connected subgraphs (HCS) clustering [21]. The HCS clustering algorithm is based on the partition of a similarity graph into all its highly connected subgraphs. More precisely, two active columns *v*_1_, *v*_2_ *∈ V* are in the same cluster if they belong to the same highly connected subgraph of *G*. As already mentioned, HCS clustering does not make any *a priori* assumptions on the number of clusters. Furthermore, it satisfies the following property: all clusters *C*_1_, *C*_2_, …, *C*_*r*_ have diameter at most 2. It guarantees that the co-evolution distance between two positions of the same cluster is at most 2*E*, where *J* (*v*_1_, *v*_2_) *< E*. Based on the coevolution distance, we construct the *similarity graph G* = (*V, E*) as follows: two active positions *v*_1_, *v*_2_ *∈ V* are connected by an edge if their *P* - or *J*-distance is smaller than the threshold *J* (*v*_1_, *v*_2_) *< E*. We then group the active positions into disjoint clusters *C*_1_, *C*_2_, …, *C*_*r*_ via HCS-clustering on the similarity graph.

### Data preparation

High-quality SARS-CoV-2 whole genome data were collected from GISAID [44]. Each sequence was individually aligned to the reference sequence collected from Wuhan, 2019 (GISAID ID: EPI_ISL_402124). A multiple sequence alignment (MSA) was produced by MAFFT [24]. We partitioned the collected sequences by week, further differentiating them by their country or region. For the scope of our analysis, we consider data from England, India, USA and SA, since these cover the regions from which the respective VOCs originate.

We consider all current VOCs Alpha, Beta, Delta (AY.3 included), Gamma, and all current VOIs Lambda, Mu. We collect the characteristic mutations of these VOCs and VOIs, including both synonymous and non-synonymous, from Outbreak.info and NextClade [33, 18]

### Analysis Protocol

We validate our framework employing curated SARS-CoV-2 sequencing data as follows: moving back in time and exclusively using the MSA, we shall reconstruct the motif complex, i.e. identify the clusters of positions corresponding to co-evolving mutations. We then take advantage of the fact that we also have an independent biological analysis including phylogenies. This in turn allows us to discuss how our clusters “fit” in the landscape of recognized VOCs and VOIs. We show that our motifs are detected weeks, if not months before the associated variants begin to be observed by other means.

We perform this analysis up to the latest genome sequence data collected from GISAID (2021-08-31) for England, India, the United States (USA), and SA. We divide a month into four weeks (1-7 for the 1st week, 8-15 for the 2nd week,16-23 for the 3rd week, and 24 after for the 4th week), and partition all aligned sequences by week. We approximate the motif complex based on nucleotide diversity and co-evolution distances, determining the 0-simplices by selecting active positions with nucleotide diversity *D*(*i*) *>* 0.1. Here nucleotide diversity accounts only for contributions from the four nucleotides {*A, U, C, G*} and neglects non-canonical symbols such as gaps and (N)s. Furthermore, if more than 30% of a column exhibits non-canonical symbols, the column will be discarded. We then approximate the 1-simplices by computing the *P* - and *J* -distance between the pairs of positions and approximate higher dimensional simplices via cluster analysis based on *k*-means an HCS clustering. For *k*-means clustering, the optimal *k* is determined by the gap statistic method [45]. For HCS clustering the similarity graph is constructed using a threshold parameter *t*, where two columns are connected by an edge in the similarity graph if their coevolution distance is *< t*. Here, we set *t*_*p*_ = 0.1 for the *P*-distance and *t*_*j*_ = 0.45 for the *J*-distance.

This results in the reconstruction of the underlying motif complex, whose high dimensional simplices correspond to the clusters of co-evolving mutations. We examine the clustering of newly emerging active positions in each week, i.e., they present in the current week but not present in the last week. We apply the criteria: a cluster of size at least five containing at least one (+)-site is considered as the signal of a potential emerging variant. Finally our findings are discussed within the context of the identified VOCs and VOIs, in particular how early can we detect blocks of positions corresponding to mutations that later were recognized to be characteristic for a VOC.

## Data Availability

The nucleotide sequences of the SARS-CoV-2 genomes used in this analysis are available, upon free registration, from the GISAID database (https://www.gisaid.org/).

## Acknowledgements

We thank Dr. Anindya Dutta and Briana Wilson for the discussion and inspiration of the work. We thank Dr. Andrew Warren for helping access SARS-CoV-2 data. Dr. Warren pointed out the importance of applying our method to VOC/VOI, monophyletic clusters and numerous references. We thank Mia Shu for helping accessing and processing GISAID data. Many thanks to Maxwell Reidys for his proofreading.

## Footnotes

1 but are not necessarily equal to

2 by means of consensus and biological analysis

3 in case of 70%-being characteristic mutations

4 finite population survival probability being exponential in population size

## References

[1] Daniel Aguilar, Baldo Oliva, and Cristina Marino Buslje. Mapping the Mutual Information Network of Enzymatic Families in the Protein Structure to Unveil Functional Features. PLOS ONE, 7(7):e41430, 2012.

[2] Daniel Aloise, Amit Deshpande, Pierre Hansen, and Preyas Popat. Np-hardness of euclidean sum-of-squares clustering. Machine learning, 75(2):245–248, 2009.

[3] T Bedford, EB Hodcroft, and RA Neher. Updated nextstrain sars cov-2 clade naming strategy. Nextstrain https://go.nature.com/3c9Riep, 2021.

[4] Anderson F. Brito, Elizaveta Semenova, Gytis Dudas, Gabriel W. Hassler, Chaney C. Kalinich, Moritz U. G. Kraemer, Sarah C. Hill, Danish Covid-19 Genome Consortium, Ester C. Sabino, Oliver G. Pybus, Christopher Dye, Samir Bhatt, Seth Flaxamn, Marc A. Suchard, Nathan D. Grubaugh, Guy Baele, and Nuno R. Faria. Global disparities in SARS-CoV-2 genomic surveillance. medRxiv: The Preprint Server for Health Sciences, 2021. DOI: 10.1101/2021.08.21.21262393.

[5] Andrei Bura, Qijun He, and Christian Reidys. Weighted Homology of Bi-Structures over Certain Discrete Valuation Rings. Mathematics, 9(7):744, 2021.

[6] Cristina Marino Buslje, Javier Santos, Jose Maria Delfino, and Morten Nielsen. Correction for phylogeny, small number of observations and data redundancy improves the identification of coevolving amino acid pairs using mutual information. Bioinformatics, 25(9):1125–1131, 2009.

[7] Thomas M. Cover and Joy A. Thomas. Elements of Information Theory (Wiley Series in Telecommunications and Signal Processing). Wiley-Interscience, New York, NY, USA, 2006.

[8] Xianding Deng, Wei Gu, Scot Federman, Louis Du Plessis, Oliver G Pybus, Nuno R Faria, Candace Wang, Guixia Yu, Brian Bushnell, Chao-Yang Pan, et al. Genomic surveillance reveals multiple introductions of sars-cov-2 into northern california. Science, 369(6503):582– 587, 2020.

[9] B Derrida and L Peliti. Evolution in a flat fitness landscape. Bulletin of Mathematical Biology, 53(3):355–382, 1991.

[10] S.D. Dunn, L.M. Wahl, and G.B. Gloor. Mutual information without the influence of phylogeny or entropy dramatically improves residue contact prediction. Bioinformatics, 24(3):333– 340, 2008.

[11] Julien Y Dutheil. Detecting coevolving positions in a molecule: why and how to account for phylogeny. Briefings in bioinformatics, 13(2):228–243, 2012.

[12] Zhenqiang Fan, Bo Yao, Yuedi Ding, Jing Zhao, Minhao Xie, and Kai Zhang. Entropy-driven amplified electrochemiluminescence biosensor for rdrp gene of sars-cov-2 detection with self-assembled dna tetrahedron scaffolds. Biosensors and Bioelectronics, 178:113015, 2021.

[13] Tom Fawcett. An introduction to roc analysis. Pattern recognition letters, 27(8):861–874, 2006.

[14] Stephen K Gire, Augustine Goba, Kristian G Andersen, Rachel SG Sealfon, Daniel J Park, Lansana Kanneh, Simbirie Jalloh, Mambu Momoh, Mohamed Fullah, Gytis Dudas, et al. Genomic surveillance elucidates ebola virus origin and transmission during the 2014 outbreak. science, 345(6202):1369–1372, 2014.

[15] Rodrigo Gouveia-Oliveira and Anders G Pedersen. Finding coevolving amino acid residues using row and column weighting of mutual information and multi-dimensional amino acid representation. Algorithms for molecular biology, 2:12, 2007.

[16] Brad Gulko and David Haussler. Using multiple alignments and phylogenetic trees to detect rna secondary structure. In Pac Symp Biocomput, pages 350–367. World Scientific, 1996.

[17] RR Gutell, A Power, GZ Hertz, EJ Putz, and GD Stormo. Identifying constraints on the higher-order structure of rna: continued development and application of comparative sequence analysis methods. Nucleic acids research, 20(21):5785–5795, 1992.

[18] James Hadfield, Colin Megill, Sidney M Bell, John Huddleston, Barney Potter, Charlton Callender, Pavel Sagulenko, Trevor Bedford, and Richard A Neher. Nextstrain: real-time tracking of pathogen evolution. Bioinformatics, 34(23):4121–4123, 2018.

[19] R. W. Hamming. Error detecting and error correcting codes. The Bell System Technical Journal, 29(2):147–160, 1950.

[20] John A Hartigan and Manchek A Wong. Ak-means clustering algorithm. Journal of the Royal Statistical Society: Series C (Applied Statistics), 28(1):100–108, 1979.

[21] Erez Hartuv and Ron Shamir. A clustering algorithm based on graph connectivity. Information Processing Letters, 76(4):175–181, 2000.

[22] Paul Jaccard. The Distribution of the Flora in the Alpine Zone.1. New Phytologist, 11(2):37– 50, 1912.

[23] Alboukadel Kassambara. Practical guide to cluster analysis in R: Unsupervised machine learning, volume 1. Sthda, 2017.

[24] Kazutaka Katoh, Kazuharu Misawa, Kei-ichi Kuma, and Takashi Miyata. Mafft: a novel method for rapid multiple sequence alignment based on fast fourier transform. Nucleic acids research, 30(14):3059–3066, 2002.

[25] Maurice G Kendall. A new measure of rank correlation. Biometrika, 30(1/2):81–93, 1938.

[26] Frank Konings, Mark D Perkins, Jens H Kuhn, Mark J Pallen, Erik J Alm, Brett N Archer, Amal Barakat, Trevor Bedford, Jinal N Bhiman, Leon Caly, et al. Sars-cov-2 variants of interest and concern naming scheme conducive for global discourse. Nature Microbiology, pages 1–3, 2021.

[27] Bette Korber, Will M Fischer, Sandrasegaram Gnanakaran, Hyejin Yoon, James Theiler, Werner Abfalterer, Nick Hengartner, Elena E Giorgi, Tanmoy Bhattacharya, Brian Foley, et al. Tracking changes in sars-cov-2 spike: evidence that d614g increases infectivity of the covid-19 virus. Cell, 182(4):812–827, 2020.

[28] S. Kullback and R. A. Leibler. On Information and Sufficiency. The Annals of Mathematical Statistics, 22(1):79–86, 1951.

[29] Sébastien Lê, Julie Josse, François Husson, et al. Factominer: an r package for multivariate analysis. Journal of statistical software, 25(1):1–18, 2008.

[30] James MacQueen et al. Some methods for classification and analysis of multivariate observations. In Proceedings of the fifth Berkeley symposium on mathematical statistics and probability, volume 1, pages 281–297. Oakland, CA, USA, 1967.

[31] Daniele Mercatelli and Federico M. Giorgi. Geographic and Genomic Distribution of SARS-CoV-2 Mutations. Frontiers in Microbiology, 11:1800, 2020.

[32] Petra Mlcochova, Steven Kemp, Mahesh Shanker Dhar, Guido Papa, Bo Meng, Isabella ATM Ferreira, Rawlings Datir, Dami A Collier, Anna Albecka, Sujeet Singh, et al. Sars-cov-2 b. 1.617. 2 delta variant replication and immune evasion. Nature, pages 1–8, 2021.

[33] Julia L. Mullen, Ginger Tsueng, Alaa Abdel Latif, Manar Alkuzweny, Narco Cano, Emily Haag, Jerry Zhou, Mark Zeller, Emory Hufbauer, Nate Matteson, Kristian G. Andersen, Chunlei Wu, Andrew I. Su, Karthik Gangavarapu, and Laura D. Hughes. https://outbreak.info, 2020. https://outbreak.info/.

[34] Jerome L Myers, Arnold Well, and Robert Frederick Lorch. Research design and statistical analysis. Routledge, 2010.

[35] Maria Pachetti, Bruna Marini, Francesca Benedetti, Fabiola Giudici, Elisabetta Mauro, Paola Storici, Claudio Masciovecchio, Silvia Angeletti, Massimo Ciccozzi, Robert C Gallo, et al. Emerging sars-cov-2 mutation hot spots include a novel rna-dependent-rna polymerase variant. Journal of translational medicine, 18(1):1–9, 2020.

[36] K Pearson. Notes on regression and inheritance in the case of two parents proceedings of the royal society of london, 58, 240–242, 1895.

[37] Jessica A Plante, Yang Liu, Jianying Liu, Hongjie Xia, Bryan A Johnson, Kumari G Lokugamage, Xianwen Zhang, Antonio E Muruato, Jing Zou, Camila R Fontes-Garfias, et al. Spike mutation d614g alters sars-cov-2 fitness. Nature, 592(7852):116–121, 2021.

[38] Prerna Priya and Asheesh Shanker. Coevolutionary forces shaping the fitness of sars-cov-2 spike glycoprotein against human receptor ace2. Infection, Genetics and Evolution, 87:104646, 2021.

[39] Andrew Rambaut, Edward C. Holmes, Áine O’Toole, Verity Hill, John T. McCrone, Christopher Ruis, Louis du Plessis, and Oliver G. Pybus. A dynamic nomenclature proposal for SARS-CoV-2 lineages to assist genomic epidemiology. Nature Microbiology, 5(11):1403–1407, 2020.

[40] Andrew Rambaut, Edward C Holmes, Áine O’Toole, Verity Hill, John T McCrone, Christopher Ruis, Louis du Plessis, and Oliver G Pybus. Addendum: A dynamic nomenclature proposal for sars-cov-2 lineages to assist genomic epidemiology. Nature Microbiology, 6(3):415– 415, 2021.

[41] Janet D Robishaw, Scott M Alter, Joshua J Solano, Richard D Shih, David L DeMets, Dennis G Maki, and Charles H Hennekens. Genomic surveillance to combat covid-19: challenges and opportunities. The Lancet Microbe, 2021.

[42] Hinrich Schütze, Christopher D Manning, and Prabhakar Raghavan. Introduction to information retrieval, volume 39. Cambridge University Press Cambridge, 2008.

[43] C. E. Shannon. A mathematical theory of communication. The Bell System Technical Journal, 27(3):379–423, 1948.

[44] Yuelong Shu and John McCauley. Gisaid: Global initiative on sharing all influenza data–from vision to reality. Eurosurveillance, 22(13):30494, 2017.

[45] Robert Tibshirani, Guenther Walther, and Trevor Hastie. Estimating the number of clusters in a data set via the gap statistic. Journal of the Royal Statistical Society: Series B (Statistical Methodology), 63(2):411–423, 2001.

[46] Elisabeth R.M. Tillier and Thomas W.H. Lui. Using multiple interdependency to separate functional from phylogenetic correlations in protein alignments. Bioinformatics, 19(6):750– 755, 2003.

[47] Rui Wang, Jiahui Chen, Kaifu Gao, Yuta Hozumi, Changchuan Yin, and Guo-Wei Wei. Analysis of SARS-CoV-2 mutations in the United States suggests presence of four substrains and novel variants. Communications Biology, 4(1):1–14, 2021.

[48] WHO. WHO announces simple, easy-to-say labels for SARS-CoV-2 variants of interest and concern, 2021. https://www.who.int.

[49] Chen-Hsiang Yeang, Jeremy FJ Darot, Harry F Noller, and David Haussler. Detecting the coevolution of biosequences—an example of rna interaction prediction. Molecular biology and evolution, 24(9):2119–2131, 2007.

