## Supplemental Materials for "Buying time: detecting Vocs in SARS-CoV-2 via co-evolutionary signals"

### Supplementary Information

The paper provides a signal detection following our analysis protocol for England, USA, India, South America SARS-CoV-2 whole-genome data. The protocol is based on k-means clustering and $P$-distance. Here, we provide additional signal detection figures using different combinations of the two clustering methods (k-means and Highly Connected Sub-graph (HCS)) and co-evolutionary measurements ($P$-distance and $J$-distance).

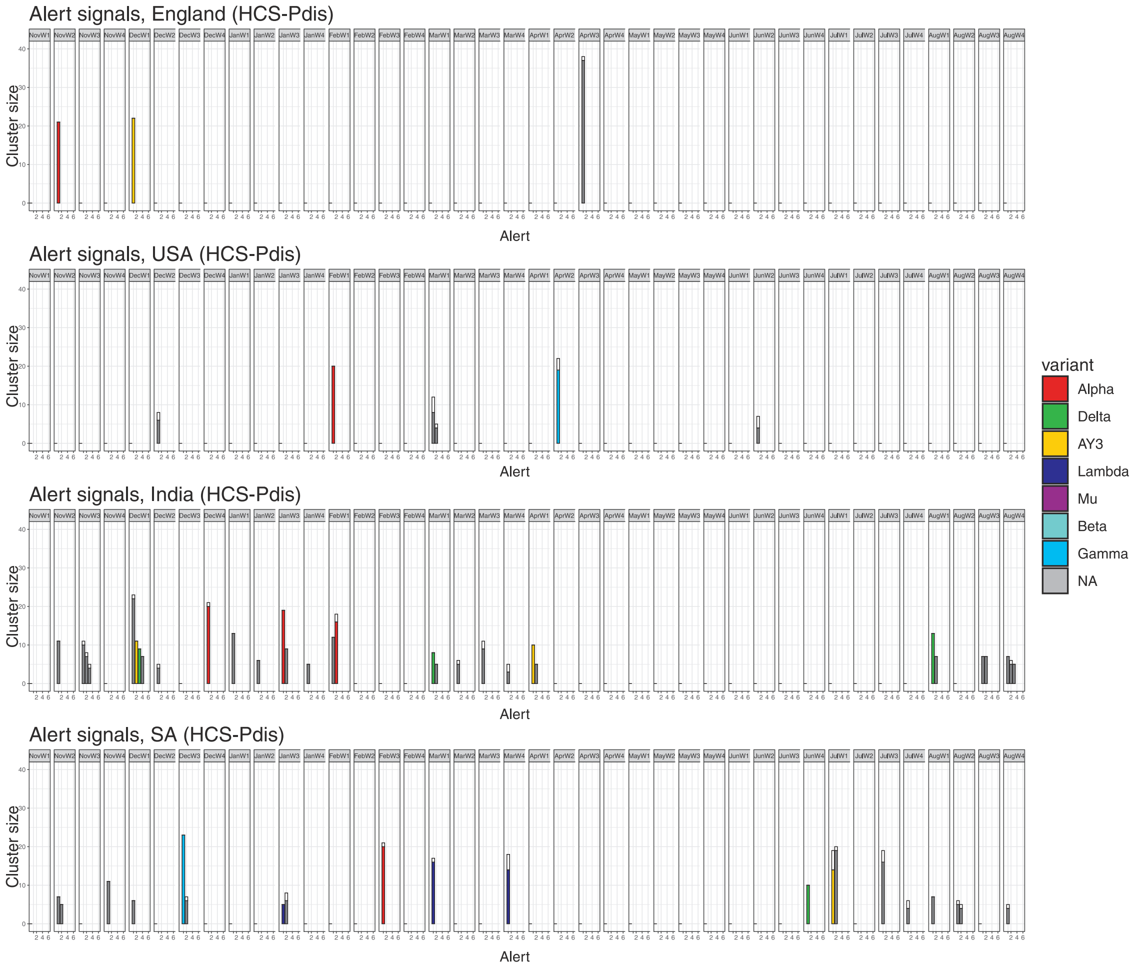


FIGURE 1. $P$-distance and HCS clustering.

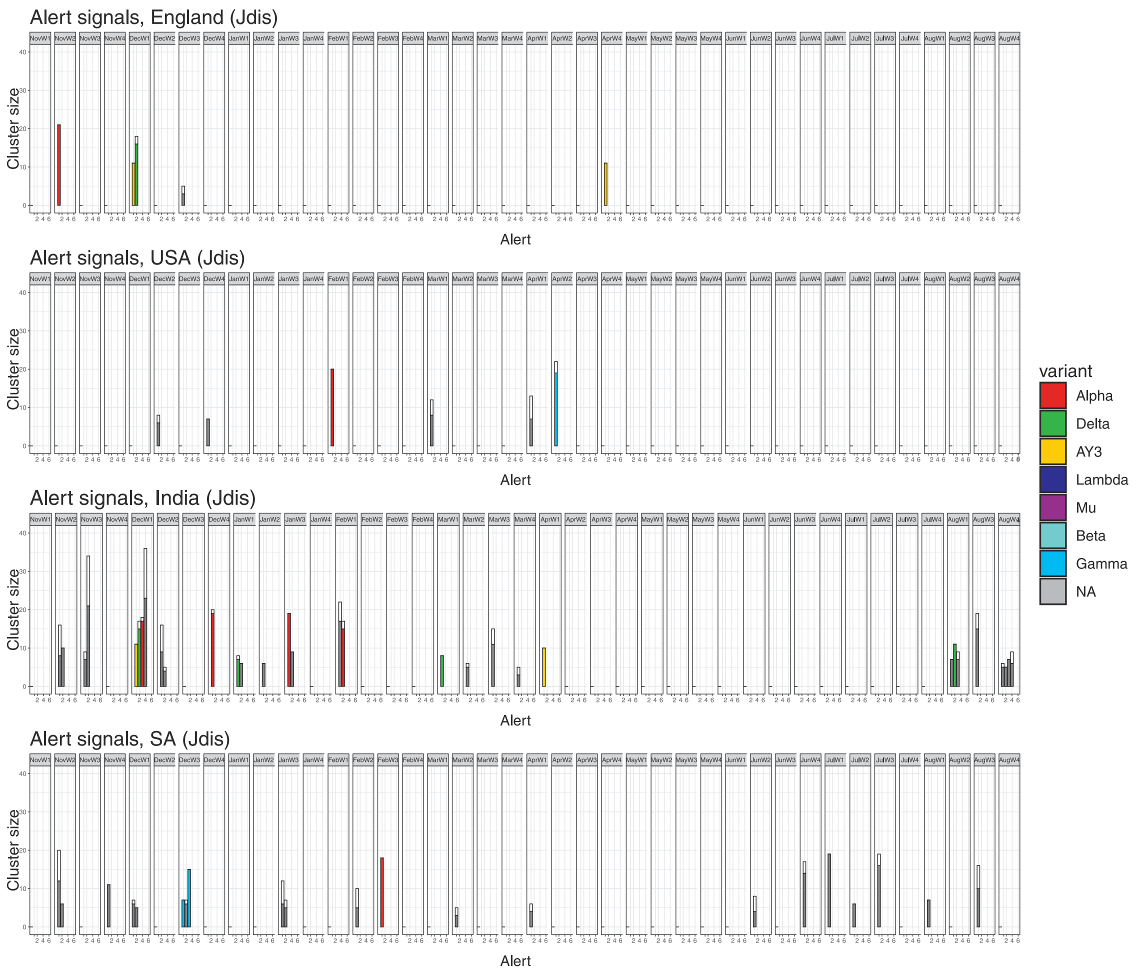


FIGURE 2. $J$-distance and k-means clustering.

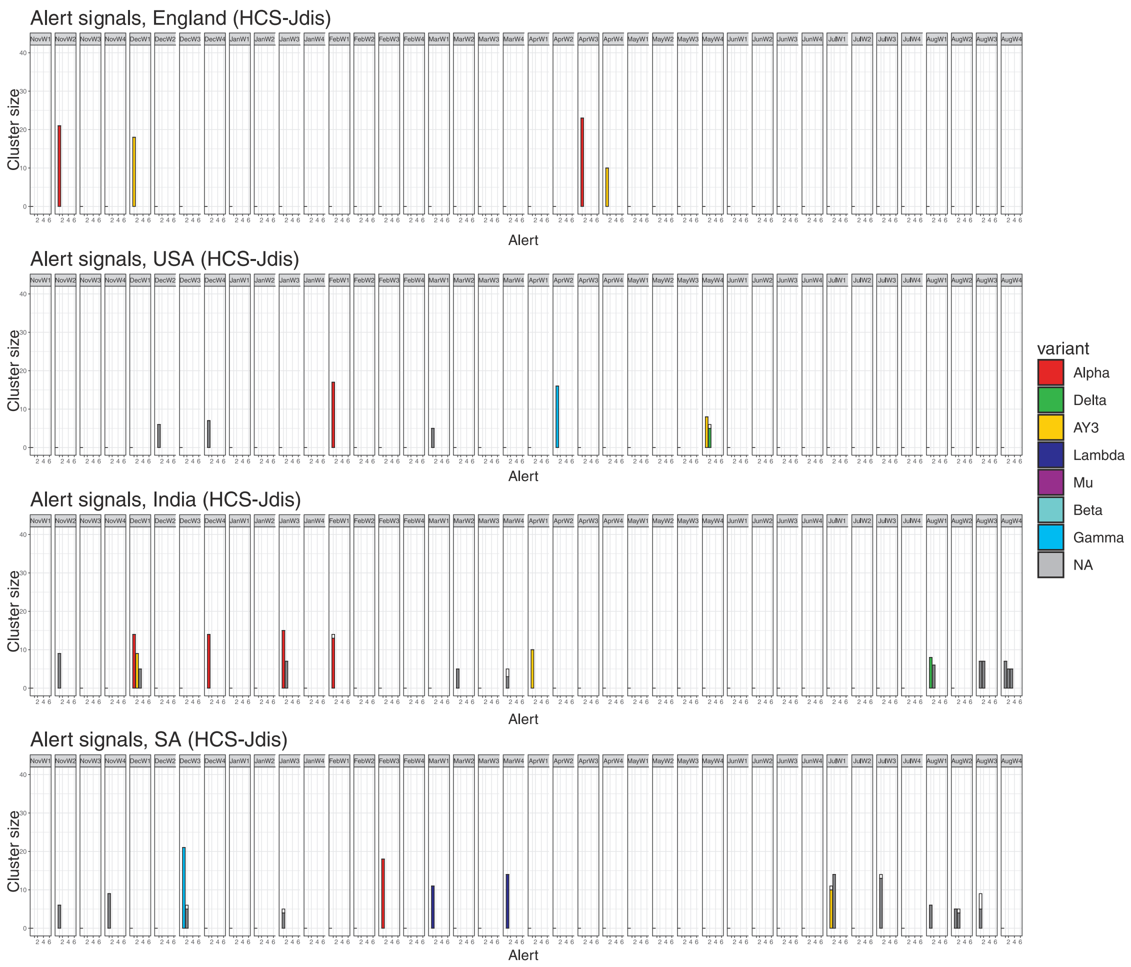


FIGURE 3. $J$-distance and HCS clustering.
